# A simple method for spray-on gene editing *in planta*

**DOI:** 10.1101/805036

**Authors:** Cara Doyle, Katie Higginbottom, Thomas A. Swift, Mark Winfield, Christopher Bellas, David Benito-Alifonso, Taryn Fletcher, M. Carmen Galan, Keith Edwards, Heather M. Whitney

## Abstract

Potential innovation in Plant research using gene-edited and genetically modified plants is currently being hindered by inefficient and costly plant transformation. We show that carbon dots formed from natural materials (quasi-spherical, <10nm nanoparticles) can act as a fast vehicle for carrying plasmids into mature plant cells, resulting in transient plant transformation in a number of important crop species with no negative impacts on photosynthesis or growth. We further show that GFP, Cas9, and gRNA introduced into wheat via foliar application (spraying on) of plasmid coated carbon dots are expressed and, in the case of Cas9, make genome edits in *SPO11* genes. Therefore, we present a protocol for spray-on gene editing that is simple, inexpensive, fast, transforms *in planta*, and is applicable to multiple crop species. We believe this technique creates many opportunities for the future of plant transformation in research and shows great promise for plant protein production systems.

## Introduction

Recent advances in plant biotech, particularly manipulation of photosynthesis, have shown the ability to obtain huge increases in plant efficiency and yield. For example, the RIPE project^1^ obtained up to a 15% increase in biomass^2^ and a ∼40% increase in productivity^3^ by reducing photoprotection latency times and by avoiding photorespiration. These examples show the true power of GM – not only could these changes increase global food security (a growing issue with our population still increasing^4,5^, and climate change conferring multiple environmental stresses^6–8^), but since these advances also increase the amount of carbon being fixed, it could have potential for also mitigating climate change^9,10^. This is important because, as noted, the effects of climate change exacerbate food insecurity further.

Plant biotechnology can also enhance food security and biomass production by improving crop resistance to herbivores, pests, and environmental stresses. Additionally, GM techniques can enhance the nutritional value of the food produced, as seen with purple tomatoes^11,12^, enhancing the lipid content of oil crops to provide an alternative to dwindling fish oil stocks^13,14^ and improving the quality of staple crops such as wheat^15,16^. However, the scope extends beyond edible compounds, as plant biotech is allowing the production of biofuels^17^, and has shown success producing pharmaceuticals^18^, including the efficient and speedy production of vaccines^19^. These advancements have been aided by new fields such as Synthetic biology, and gene editing tools becoming more versatile and useable.

However, there is currently a significant bottleneck^20^ limiting the potential application of these ideas and advances, and that is the cost, both in time and resources, of current plant transformation methods. All plant transformation must currently utilise either *Agrobacterium tumefaciens*^21^, biolistics^22^, or regeneration from PEG transformed protoplasts^23^, as a vehicle to introduce DNA, regardless of whether the changes are transient (non-heritable) or stable (inheritable). This means that not only are established advances such as GM species and cultivar limited, but new technologies such as gene editing^24,25^ suffer the same bottleneck as these delivery methods are still required. Furthermore, even in species and cultivars where plant transformation is possible, the process is expensive, slow, requires significant resources in terms of facilities and expertise, is frequently inefficient^26^, and damages the plant genome^27,28^.

Carbon nanomaterials have found multiple uses since their discovery in 2004 due to their varied shapes (including nanosheets^29^, nanotubes^30,31^, nanodots) and sizes. Carbon dots (CDs) can be functionalised by PEG diamines^32^, allowing whole plasmids to electrostatically interact with them, and thus can act as a vehicle to carry the plasmid into plant cells. They have the advantage of occurring naturally or being made from natural non-toxic materials^33^. CDs are easy, fast, and inexpensive to make, require little equipment to do so, and, due to the variety of application routes offered, presents itself as a simple way to obtain GM or gene-edited plants across model organisms, crop plants, and orphan crops (plants notoriously recalcitrant to transformation). We show transformation is obtained with no decreases in photosynthetic efficiency^34^ and no growth defects.

## Results

### Synthesis of CD-plasmid complexes

The synthesis of the CD-plasmid complexes is outlined in Fig. 1. The PEG functionalised CDs were prepared by adapting the previously reported synthesis^32^ and were characterised by NMR, absorbance spectroscopy and fluorescence spectroscopy (Figs. S1–S4). The formation of a complex between the plasmid and CD was demonstrated by dynamic light scattering (DLS), Fig. S5. Further details of synthesis and characterisation are given in the supporting information.

**Figure 1.**
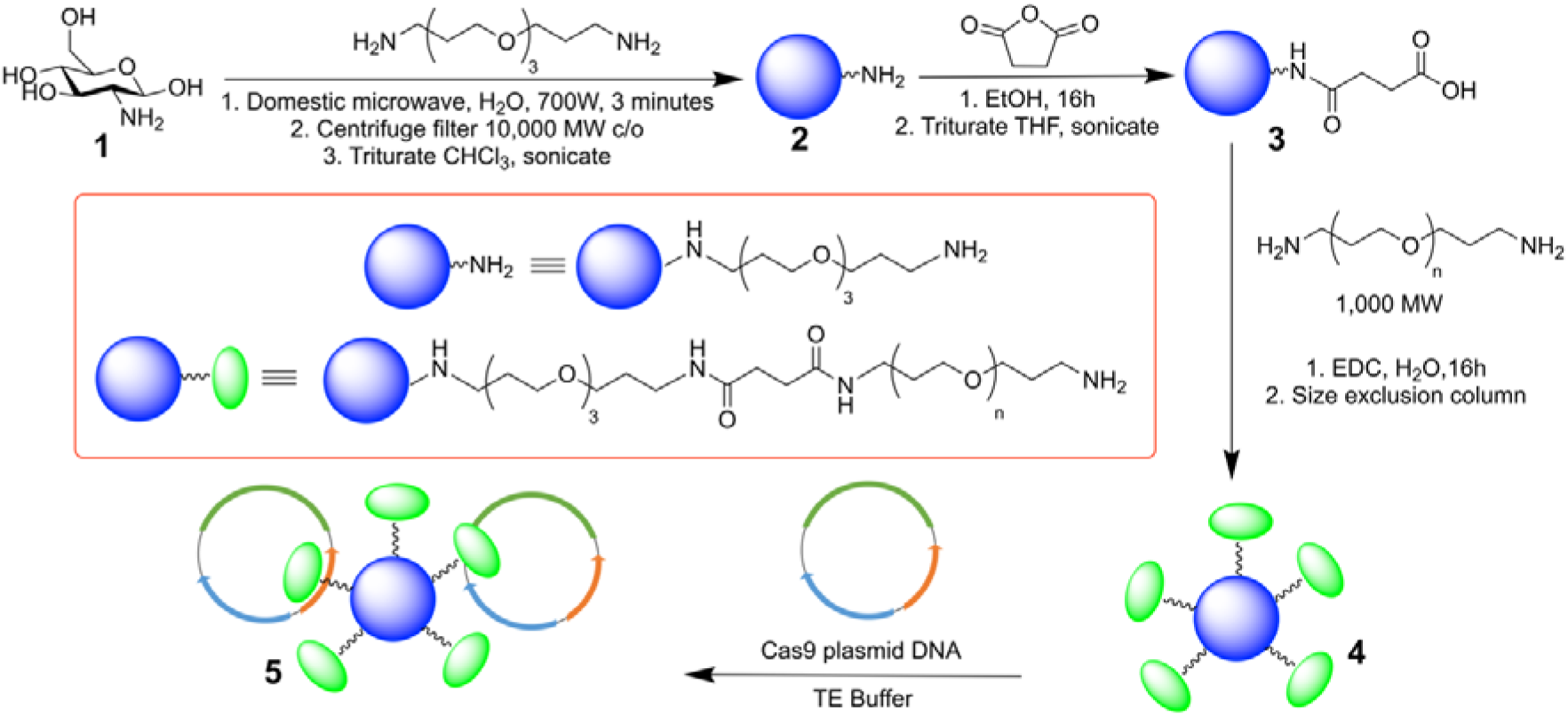
CD-plasmid complex production. Microwave-assisted reaction of 4,7,10-trioxa-1,13-tridecanediamine (TTDDA) and glucosamine hydrochloride (**1)** in water yielded amine-functionalised CDs (**2)**, which were then carboxylic acid-functionalised (**3**) via the ring-opening of succinic anhydride prior to N-(3-Dimethylaminopropyl)-N′-ethylcarbodiimide hydrochloride (EDC) mediated amide coupling with 1,000 molecular weight poly (ethylene glycol) diamine to form PEG-CDs (**4)**. The positive charge of the exposed amines on (**4**) are then used to form a complex (**5**) with partially negatively charged plasmid DNA.

### Uptake of CDs

A range of plant species readily uptake CDs delivered via foliar spray, using a simple plant mister (Fig. 2). This does not require any mechanical damage to the leaf to create routes of entry for the CDs prior to spray on application. The CDs are easily trackable within the plant without the need for a reporter gene such as GFP due to their innate fluorescence at 475nm. We saw this fluorescence in wheat after spraying, and conversely did not see this fluorescence in control plants that were not treated with CDs (Figs. 2**A** and 2**B**). We also saw that, in wheat, the CDs did not co-localise to the chloroplasts, but were generally diffuse within the somatic cells (Figs. 2**C**, 2**D** and 2**E**.)

**Figure 2.**
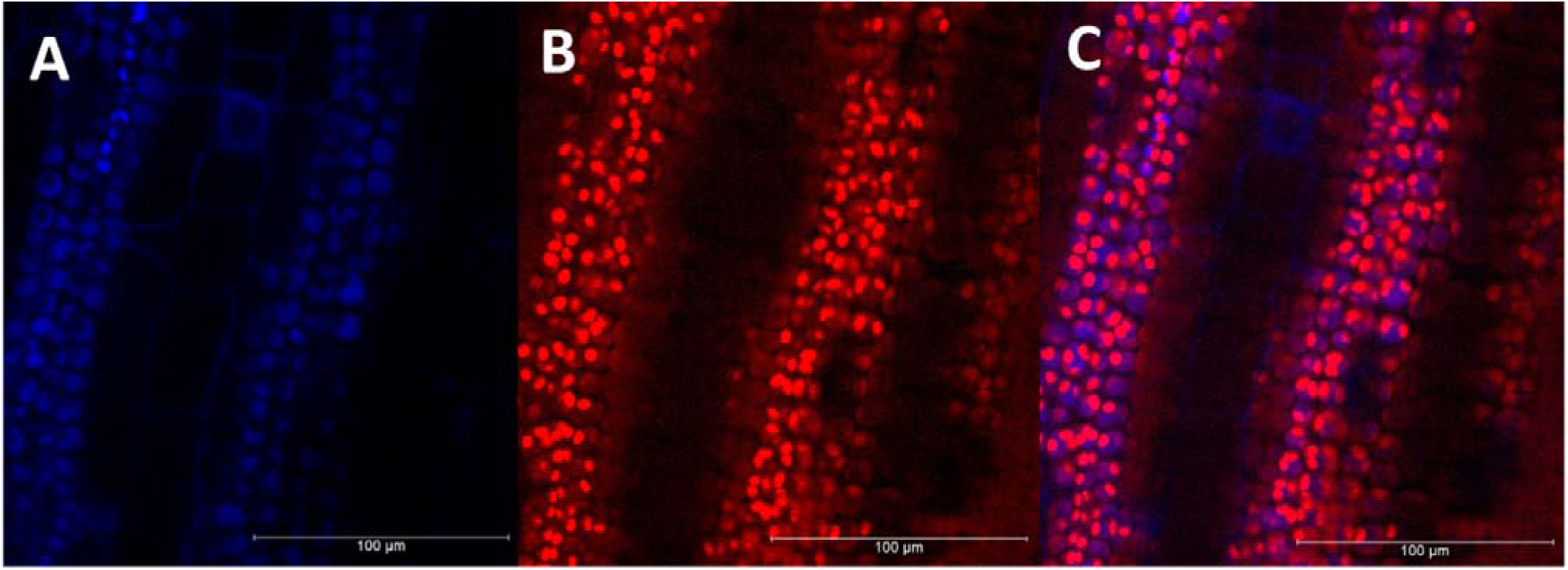
Uptake of fluorescent CDs. **(A)** CD fluorescence in wheat leaves after application via spraying (emission 475nm); **(B)** chlorophyll fluorescence of **(A)** (emission 644-713nm); **(C)** overlay of **(A)** and **(B)**.

### Nuclear expression of GFP

CD-plasmid complexes (see methods) containing a green fluorescent protein (GFP) gene with nuclear localisation sequences (NLS) were sprayed onto a range of plant species. The CDs successfully delivered the plasmid to plant cells and the GFP protein targeted the nucleus. Nuclear localised GFP fluorescence was not seen when either the CDs or the plasmid DNA were absent (Fig. 3). We saw nuclear expression of GFP in a range of species, including the important cereal crops, wheat and maize (Fig. 3), barley, and the orphan crop sorghum. The transformation efficiency of wheat was 27.74%.

**Figure 3:**
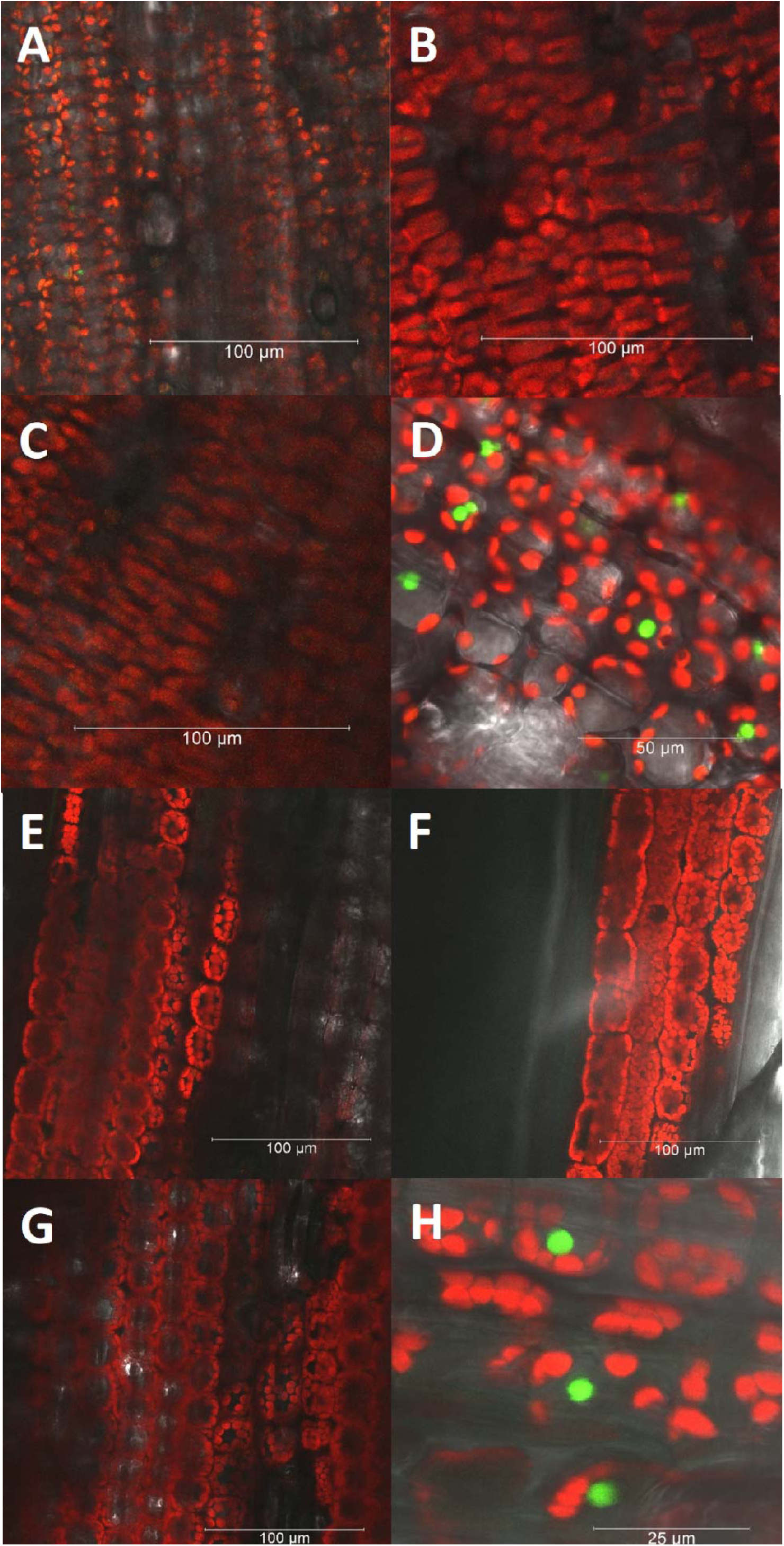
Nuclear expression of GFP reporter gene in wheat and maize after spray on application of carbon nanodots carrying a Cas9-GFP plasmid. **(A, E)** Leaf tissue sprayed with CDs carrying no plasmid show chlorophyll fluorescence (emission 644-713nm) but no GFP reporter gene expression; **(B, F)** leaf tissues sprayed with buffer containing plasmid but no CDs (no vehicle present to take DNA into cells) also do not express GFP; **(C, G)** leaf tissue sprayed only with TE buffer do not express GFP; **(D, H)** the application of the buffer solution containing both CDs and plasmid results in reporter gene expression (nuclear localised GFP fluorescence - emission 500-540nm). All images were taken 5 days after final treatment application. **(A – D)** maize, **(E – H)** wheat.

### Spray-on gene editing in wheat

The plasmid also carried the Cas9 gene and guide RNAs (gRNA) to make a deletion. The gRNA targeted Cas9 to two regions in the wheat *SPO11* genes, ~250bp apart, resulting in the deletion of the sequence in between. Please note that wheat is hexaploid and so has 6 copies of the gene. Following spray on application of CDs carrying the Cas9 plasmid to wheat, edits to *SPO11* were observed resulting in a PCR product smaller (c. 230 bp) than that from the unedited gene (Fig. 4). Presence of the gene edits was also confirmed by DNA sequencing the band shown in Fig. 4**A**, and can be found in the supporting information (Fig. S8). Appropriate controls were performed, including a positive control using standard transformation of wheat protoplasts, shown here in Fig 4**B**, and further in Fig. S9.

**Figure 4.**
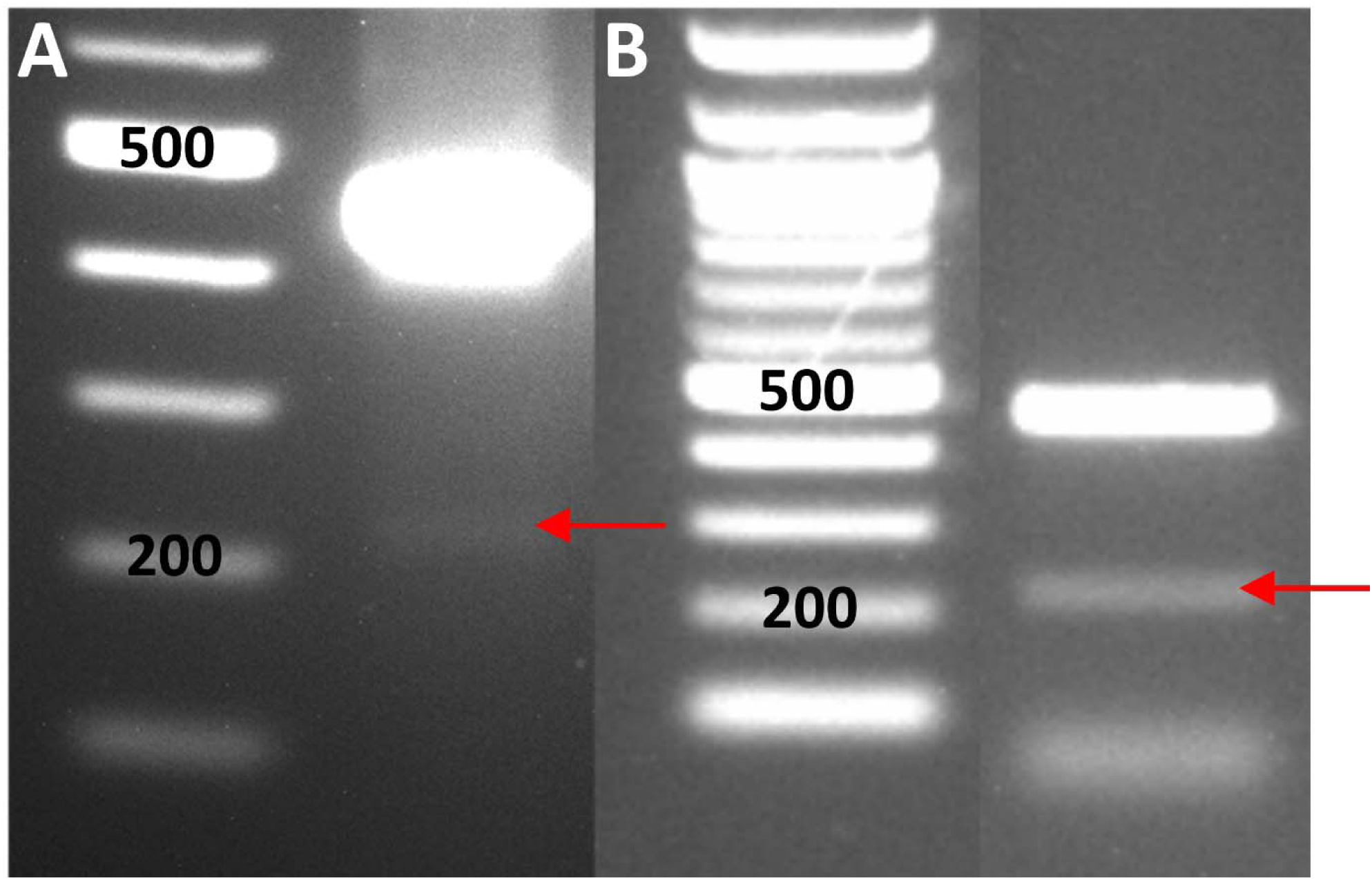
Gene editing. **(A), (B)** Gene editing resulting in bands at c.230bp, obtained by **(A)**; CD sprayed leaf material, and **(B)** standard PEG-based transformation of protoplasts. Gene editing bands are indicated by red arrows. **(A)**, Lane 1, MW ladder (bp), Lane 2, DNA extracted from a wheat leaf, after CD spraying. **(B)**, Lane 1, MW ladder (bp), Lane 2 (non-contiguous), DNA extracted from protoplasts, transformed with the same plasmid, using traditional methods.

### Versatility of application

The CDs are able to carry the plasmid into plant cells, resulting in GFP expression via multiple application routes. The GFP expression seen using the CDs as the transformation vehicle is equivalent to that obtained when using an established transformation method, protoplast transformation, using a GFP expression plasmid. (Fig. 5).

**Figure 5.**
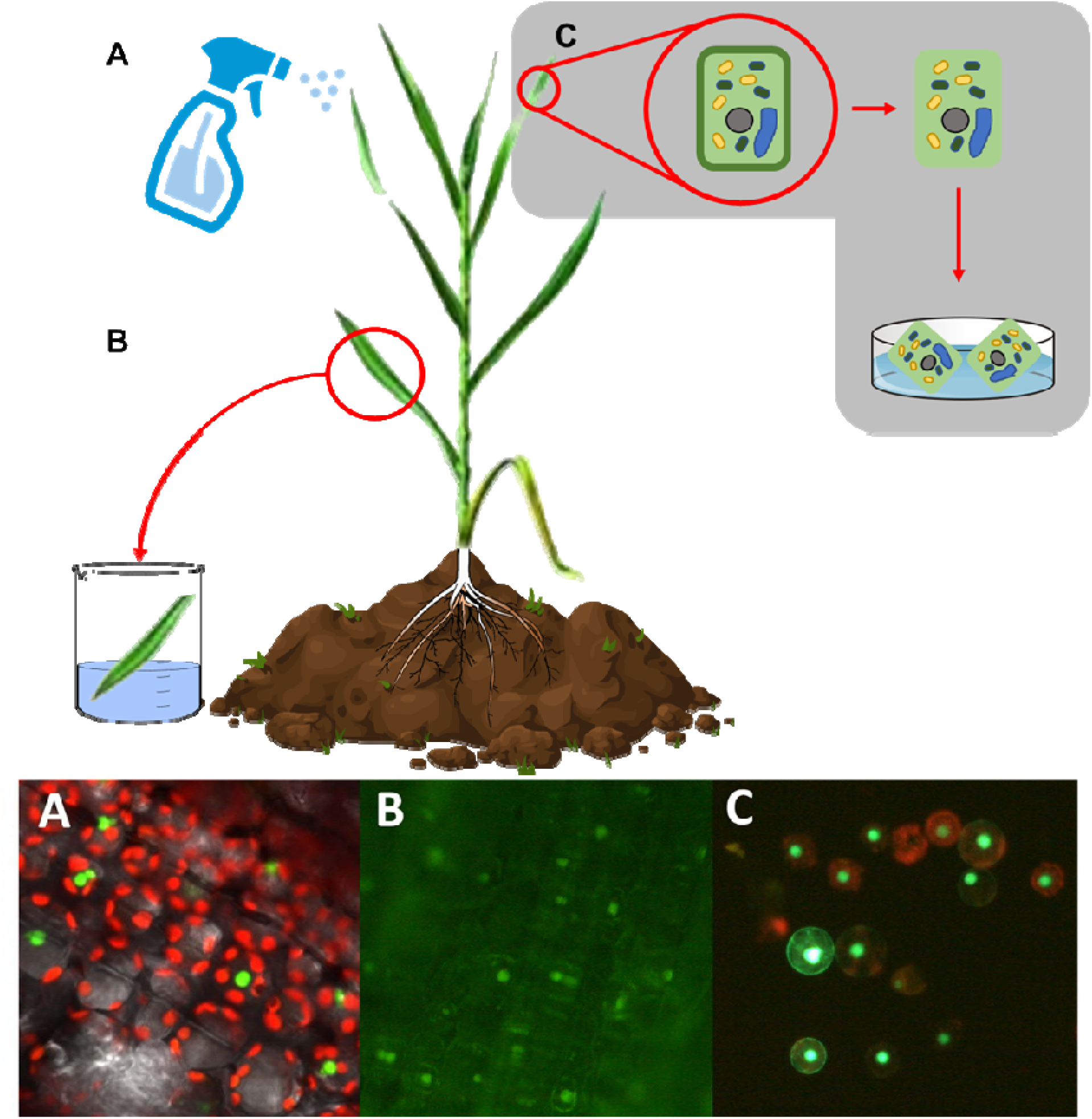
Two CD application methods that produce transformants. **(A)**, CD/plasmid complex is sprayed onto plants, **(B)**, leaves of plant dipped into CD/plasmid complex, **(C)**, positive control, PEG mediated protoplast transformation (see methods). Nuclear localised GFP fluorescence - emission 500-540nm, chlorophyll fluorescence - emission 644-713nm.

## Discussion

We show that CD–plasmid nanocomplexes can act as a delivery vehicle by which plasmids can be carried into plant somatic cells, allowing transient transformation. This new system has many advantages, including its versatility in, and ease of application. The CDs are readily taken up into mature plant tissue simply through a spray on application that requires no surface damage to the plant. Thus, the CD technique will avoid the negative side-effect of creating routes of entry for DNA that could then be utilised by plant pathogens (viruses, fungi or bacteria) to enter into plant tissue. Additionally, the CDs are non-toxic, do not cause growth defects or decrease photosynthetic efficiency even after extensive repeat application (Figs. S6 and S7), and present minimal risk to operators^35^. We have shown the expression of nuclear-localised GFP in a range of plant species including the important cereal crops, wheat, maize and barley and the orphan crop, sorghum. Importantly, we have shown that by spraying CD-plasmid nanocomplexes onto intact leaves we are able to obtain editing to the genome.

Genetically modified/edited plants are subject to strict regulation to prevent the escape of edited plants and genes which could potentially have a detrimental effect on natural ecosystems. Consequently, we acknowledge that the very simplicity and cost-effective nature of this new method means it has the potential to be misused. However, we believe that the positive benefits to the field of plant transformation far outweigh any potential negatives. The development of a plant transformation protocol that would be effective in many plant species and cultivars, including orphan crops, could bring huge benefits^36^. We believe, given our success with several important cereal crops, and with the orphan crop sorghum, which has proved to be recalcitrant to transformation, that this CD based method, if optimised, has this potential. The simplicity, versatility, and non-toxicity of this technique may also open up the use of plant transformation in situations and plant species where current cost, facilities and lack of training may limit potential.

We show here the first example of spray on gene editing. We can currently make transient gene edits in the somatic cells of plants using this CD method, but we expect refinement and optimisation will permit us to make stably gene edited lines by targeting the cells of the plant germline. However, even for use in transient transformation, this approach shows great promise: it could be used to transiently silence or increase gene expression which would be particularly useful for plant developmental research. Transient transformation is also becoming increasingly important in the production of recombinant pharmaceutical and industrial proteins *in planta* over traditional mammalian and bacterial production systems due to lower costs, minimal infection risks, and greater potential for large-scale expansion^37^. Examples of this include a multitude of vaccines (including H1N1 influenza, non-Hodgkin’s lymphoma, and Newcastle disease in poultry), pharmaceutical proteins (including interleukins, Gastric lipase, and human insulin), and industrial enzymes (including, Trypsin, lysozyme, and avidin)^38^. Transient expression allows the plant to focus on growing to maturity quickly without using resources on protein production, and then switching focus to producing the protein of interest once it is fully developed. The versatility of this method would readily allow automation via use of mist sprayers, sprinklers, or automated dipping machines as already exist for floral dip procedures, allowing ease of use for industrial/pharmaceutical protein production and research alike. Automation could allow transient transformations to be generated on an as of yet unseen scale, with previously unseen speed and efficiency. Genetic modification/editing of plants has, due to its limiting factors, struggled to find easy use across the wider scientific community and the transformation technology presented here, with its multiple benefits over traditional plant transformation methods, could allow greater uptake of plant transformation research globally.

## Methods

### Production of CDs

Core-CD synthesis and PEG functionalisation were performed using a modified version of the synthesis previously reported by Swift *et al.*,^32^. Full details of the synthesis and characterisation are given in the SI.

### Plant Material

*Triticum aestivum L.* cultivar USU-Apogee, *Hordeum vulgare L.* and *Zea mays L.* were grown in the following conditions: day temperature 20°C; night temperature 20°C; supplementary lighting duration 16 hours (5:00–21:00); thiacloprid added at 0.4g/l soil; watered daily. *Sorghum bicolor* cultivar Serredo was grown in the following conditions: day temperature 28°C; night temperature 28°C; supplementary lighting duration 16 hours (5:00–21:00); thiacloprid added at 0.4g/l soil; watered daily.

### Plasmid

The structure of the pCas9-GFP plasmid is reported in Zhang *et al*., 2019^25^.

### CD spray application

CD-DNA treatment (0.00425 X TE buffer, 0.21mg/ml PEG functionalised CDs, 0.0085mg/ml pCas9-GFP plasmid DNA, in 20ml distilled water), TE control (0.00425 X TE buffer in 20ml distilled water), TE + CD control (0.00425 X TE buffer, 0.21mg/ml PEG CDs in 20ml distilled water) and TE + DNA control (0.00425 X TE buffer, 0.0085mg/ml pCas9-GFP plasmid DNA in 20ml distilled water) were mixed as described and aliquoted into 100ml travel spray bottles. Plants were sprayed twice a day (09:00 and 15:00) for five days from ~10cm away until the plants were dripping wet. The first two true leaves (x 3 replicates) were harvested an hour after the last spray each day and immediately snap frozen in liquid nitrogen. These were stored at – 80°C until required for PCR. Sprayed plants were left for 5 days to allow production of GFP before imaging. Transformation efficiency was calculated by image analysis in ImageJ, comparing the number of cells expressing nuclear localised GFP to total number of cells visible.

### Protoplast isolation and transformation

Protoplast isolation and transformation were performed as in Shan *et al.*, 2014^39^, with the following modifications: Wheat seedlings were germinated and grown at 25°C in the dark for 9 days. All centrifugation steps were carried out at 80g rather than 250g. After isolation of protoplasts they were resuspended at a concentration of 1 × 10^6^ cells per ml rather than 2.5 × 10^6^. To 100μl of protoplasts (5 × 10^5^ cells) 10μl of a standard nuclear localised GFP expression plasmid (1μg per μl) was added and mixed gently. Transformed protoplasts were observed after 24–48 hours using a Leica DM200 and Leica MC120HD detector.

### Seed transformation

Surface sterilised seeds (*Arabidopsis thaliana* and *Triticum aestivum*) were placed in 50ml falcon tubes with 25ml liquid MS 4.4 g/l as recommend by the manufacturer and incubated at 22°C, shaking at 120 rpm with constant light for 24 hours.

The MS was removed, and the seeds were separated into four 50ml falcon tubes. 25ml liquid MS was added to each with the conditions and solution specifications as above. However, the amounts of CDs, DNA and TE buffer pH 8.0 were increased to 85µl to improve uptake in a more dilute end solution. The tubes were incubated in the same manner for another 24 hours. The seeds were washed with distilled water three times. This method was adapted from Feldmann and Marks, 1987^40^.

Seeds were pipetted onto MS30 plates, made using 4.4 g/l MS, 30 g/l sucrose and 8 g/l agar pH 5.8. The plates were sealed with parafilm and incubated in a Micro Clima-series economic lux chamber (Snijders Labs, Tilburg, Netherlands) with day cycles of 25°C for 16 hours and night cycles at 22°C for 8 hours. The plates were placed upright to allow stem and root extraction from the surface of the agar.

### DNA Extraction

Leaf tissue or protoplasts were spun using a Benchmark MC-12 (Thomas Scientific) at max speed for 2 minutes to form a pellet, then supernatant removed. 600μl of heated extraction (0.1M Tris-HCl, pH 7.5, 0.05M EDTA, 1.25% SDS) buffer was added to resuspend the pellet, which was incubated at 55°C for 20 mins. Suspension was incubated at 55°C for 20 mins. Tubes were placed on ice for 5 mins. 300μl of cold 6M ammonium acetate was added and shook vigorously then placed on ice for 15 mins. Tubes were spun for 15 mins at max speed to precipitate proteins and tissue. The supernatant was recovered, and DNA was precipitated using a standard iso-propanol procedure. The pellet was washed with 70% ethanol and then resuspended in 100μl TE buffer.

### Confocal microscopy

A Leica SP5-AOBS confocal laser scanning microscope attached to a Leica DM I6000 inverted epifluorescence Microscope was used with the following settings to image chlorophyll, GFP and CD fluorescence *in vivo*: GFP 488nm excitation, 500-540nm emission; Chlorophyll 514nm excitation, 644-713nm emission; CDs 405nm excitation, 415-470nm emission.

### PCR

To enhance PCR detection of edited *SPO11*, we used a restriction digest to first reduce the frequency of unedited copies from the DNA extract. DNA was incubated for 1 hour at 37°C with EcoRI. PCR amplification was then carried out in a 25μl reaction volume using 2X Hot Start Taq 2X Master Mix (NEB) according to the manufacturer’s recommendations and using the following primers:

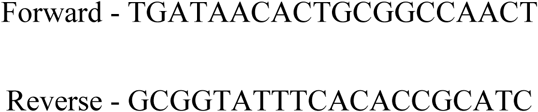

from Eurofins Genomics (Ebersberg, Germany). The amplification programme was as follow: 95°C for 5 minutes, then 40 cycles of 95°C for 30 seconds, 58°C for 30 seconds, 72°C for 60 seconds, then 72°C for 10 minutes, then held at 10°C. Samples were visualised on a 1.5% w v^−1^ agarose gel.

## Supporting information

Safety guidance

## Acknowledgements

We would like to thank Tom Pitman, Anna Lim and Alanna Kelly (University of Bristol) for growing the plants. This research was supported by the Biotechnology and Biological Sciences Research Council-funded South West Biosciences Doctoral Training Partnership BB/M009122/1. (KH); EPSRC Doctoral Prize fellowship EP/R513179/1 (TAS); EPSRC CAF EP/J002542/1 and ERC-COG:648239 (MCG), the Bristol Centre for Agricultural Innovation and the Wolfson Foundation.

## Author Contributions

TAS, HMW and MCG conceived and designed the experiments, KH, CD, TAS, MW, DBA, CB, TF collected the data, KH, CD, TAS, MW performed data analysis, KH, TAS, MW, CD wrote the manuscript, KH, CD, TAS, MW, HMW, MCG, KE proof-read the manuscript.

## Author Interests

The authors declare no competing interests.

## Supplementary

### Synthesis

Chemicals were purchased and used without further purification. Extracts were concentrated under reduced pressure using both a Büchi rotary evaporator at a pressure of 15mmHg (diaphragm pump) at 40°C. A single batch of CDs were used for all experiments.

Glucosamine HCl (1.00 g, **1**) and 4,7,10-trioxa-1,13-tridecanediamine (TTDDA) (1.35 ml) were mixed with 20ml double-distilled water (ddH_2_O). This mixture was then heated in a domestic microwave (Tesco Homebrand, 3 mins, 700 Watts.)

The resulting brown oil (**2**) was washed and sonicated with CHCl_3_ several times, discarding the supernatant, until the supernatant is clear. The brown oil was then dissolved in 20ml ddH_2_O and centrifuged through GE Healthcare Life Sciences VIVASPIN 20 with a 10,000 Da MWCO filter (5000 rpm, 1 hour) to form the core-CDs (**2**).

To form the acid decorated CDs (**3**) the core CDs (**2**) were dissolved in methanol to a concentration of 10mg/ml and sonicated. This solution was then passed through a 200nm syringe filter and mixed with 0.5 equivalence by weight of succinic anhydride. The solution was subsequently stirred vigorously overnight. The resulting solution was then reduced and the brown oil washed and sonicated with tetrahydrofuran several times, discarding the supernatant, until the supernatant was clear. The resulting brown oil was then dissolved in methanol, reduced and weighed. The brown oil was then dissolved in ddH_2_O to a concentration of 10mg/ml and sonicated for 5 minutes. This yields the acid decorated CDs (**3**).

To form the PEG functionalised CDs (**4**) 1000 molecular weight poly (ethylene glycol) diamine (PEGDA) is bonded to the acid decorated CDs (**3**) by an amide bond. To achieve this the previous solution of acid decorated CDs (**3**) was passed through a 200nm syringe filter and mixed with 2 equivalents by weight of N-(3-Dimethylaminopropyl)-N’-ethylcarbodiimide (EDC) and 10 equivalents by weight of PEGDA. PEG conjugation was performed with an excess of PEGDA to ensure all the acid groups reacted. This solution was then stirred vigorously overnight. The sample was then passed through a 200nm syringe filter. This was then purified by size-exclusion chromatography (Sephadex G-10, Sigma). The PEG-CDs were identified as a brown band on the column and were identified by both fluorescence and absorbance spectroscopy, both tails of the CD band were discarded, particularly the lower molecular weight tail as this may contain non-functionalised CDs. The resulting fraction was freeze-dried, weighed and suspended in ddH_2_O.

For storage, the PEG-CDs (**4**) were dissolved in ddH_2_O and kept at 4°C to prevent aggregation.

### Nuclear magnetic resonance (NMR)

The ^1^H 500MHz NMR spectra of the PEG-CDs are shown below in figure S1. All samples were dissolved at 1mg/ml in 0.8ml D_2_O using Norrell Select Series 7" tubes (S-5-500-7). All spectra were taken on a Bruker Advance III HD 500 Cryo. NMR chemical shifts are quoted in parts per million (ppm) and referenced to the residual solvent peak (D_2_O: δ= 4.70). Coupling constants (J) given in Hertz. Multiplicities are abbreviated as: s (singlet), d (doublet), t (triplet), q (quartet), p (pentet) and m (multiplet).

**Figure S1:**
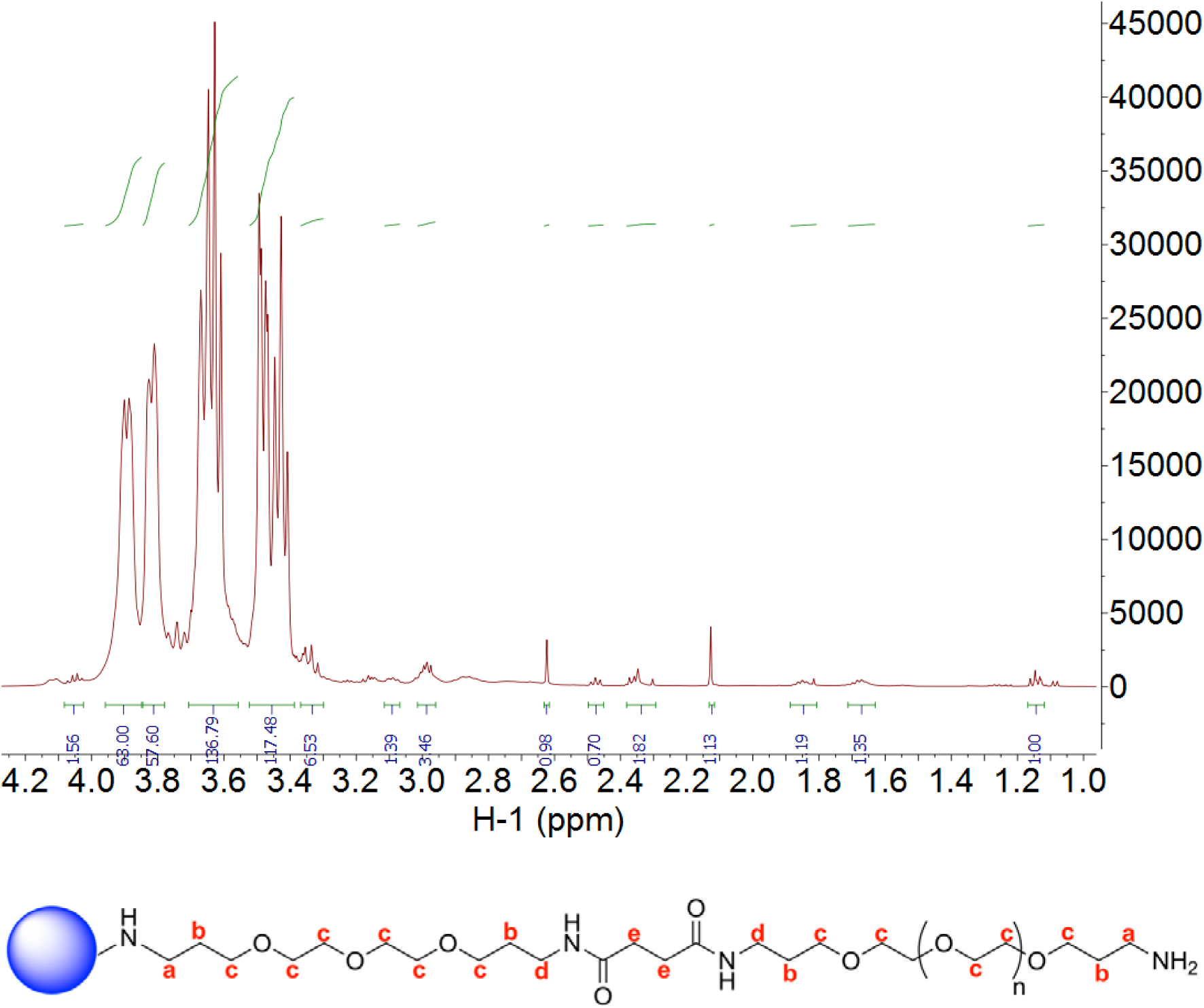
The ^1^H NMR spectra of the PEG-CDs. δ: 3.89 (m, **c**), 3.85 – 3.78 (m, **c**), 3.71 – 3.56 (m, **c**), 3.52 – 3.39 (m, **c**), 3.37 – 3.30 (m, **c**), 3.11 – 3.07 (m, **d**), 2.99 (m, **d**), 2.48 (m, **e**), 2.38 – 2.29 (m, **a**), 1.89 – 1.81 (m, **b**), 1.67 (m, **b**).

**Figure S2:**
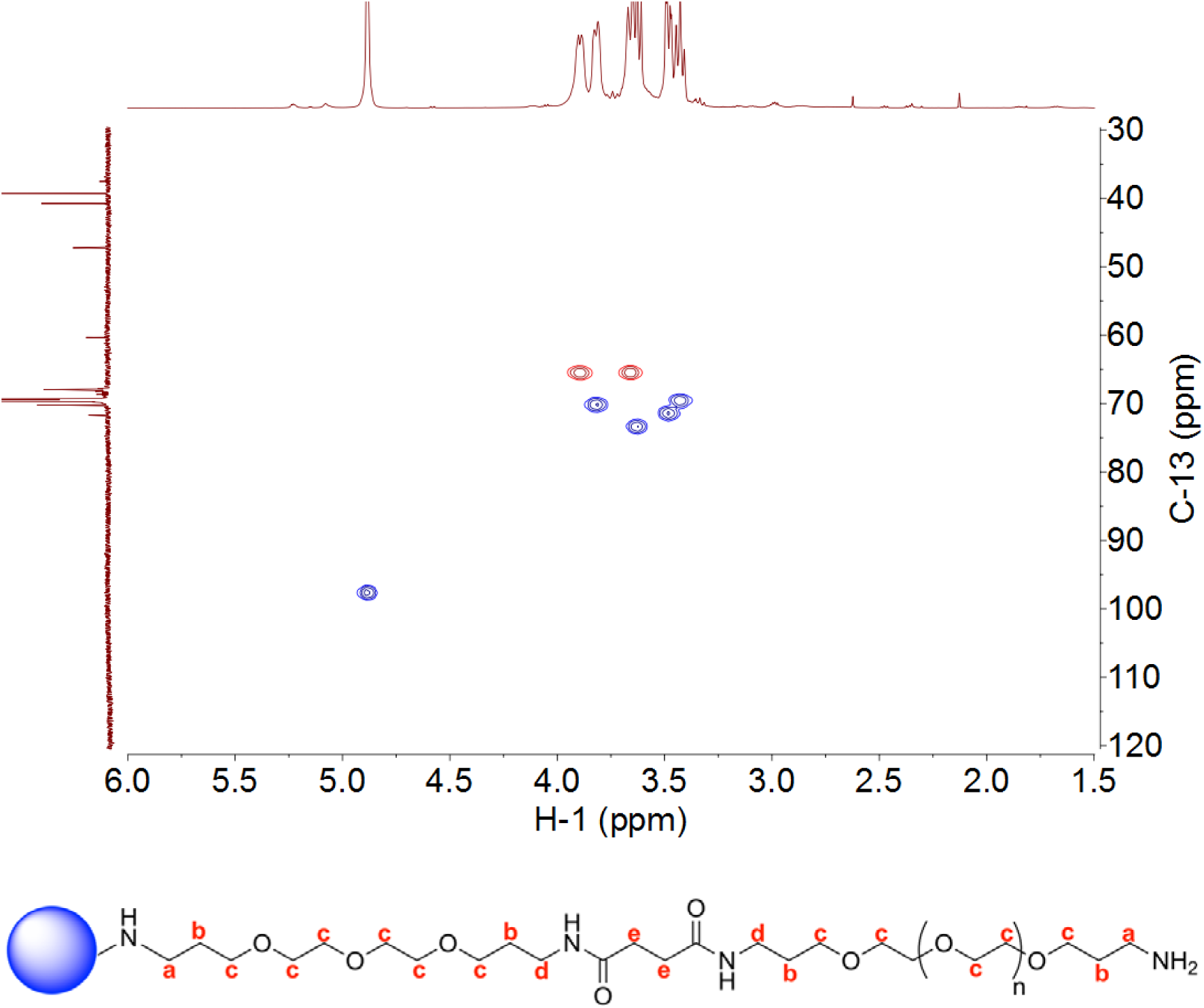
The ^1^H-^13^C HSQC spectra of the PEG-CDs.The HSQC is dominated by the signal from the **c** protons δ: 3.89 (m, **c**), 3.85 – 3.78 (m, **c**), 3.71 – 3.56 (m, **c**), 3.52 – 3.39 (m, **c**), 3.37 – 3.30 (m, **c**), forcing the other peaks below detection except the solvent, D_2_O δ=4.70

### Absorbance spectroscopy

The ultraviolet-visible absorption spectra of GdCNPs were recorded using a Cary UV-Visible 50 spectrophotometer. The absorption spectra are dominated by shoulders at ~230 and <200nm as well as broad absorption at <500nm.

**Figure S3:**
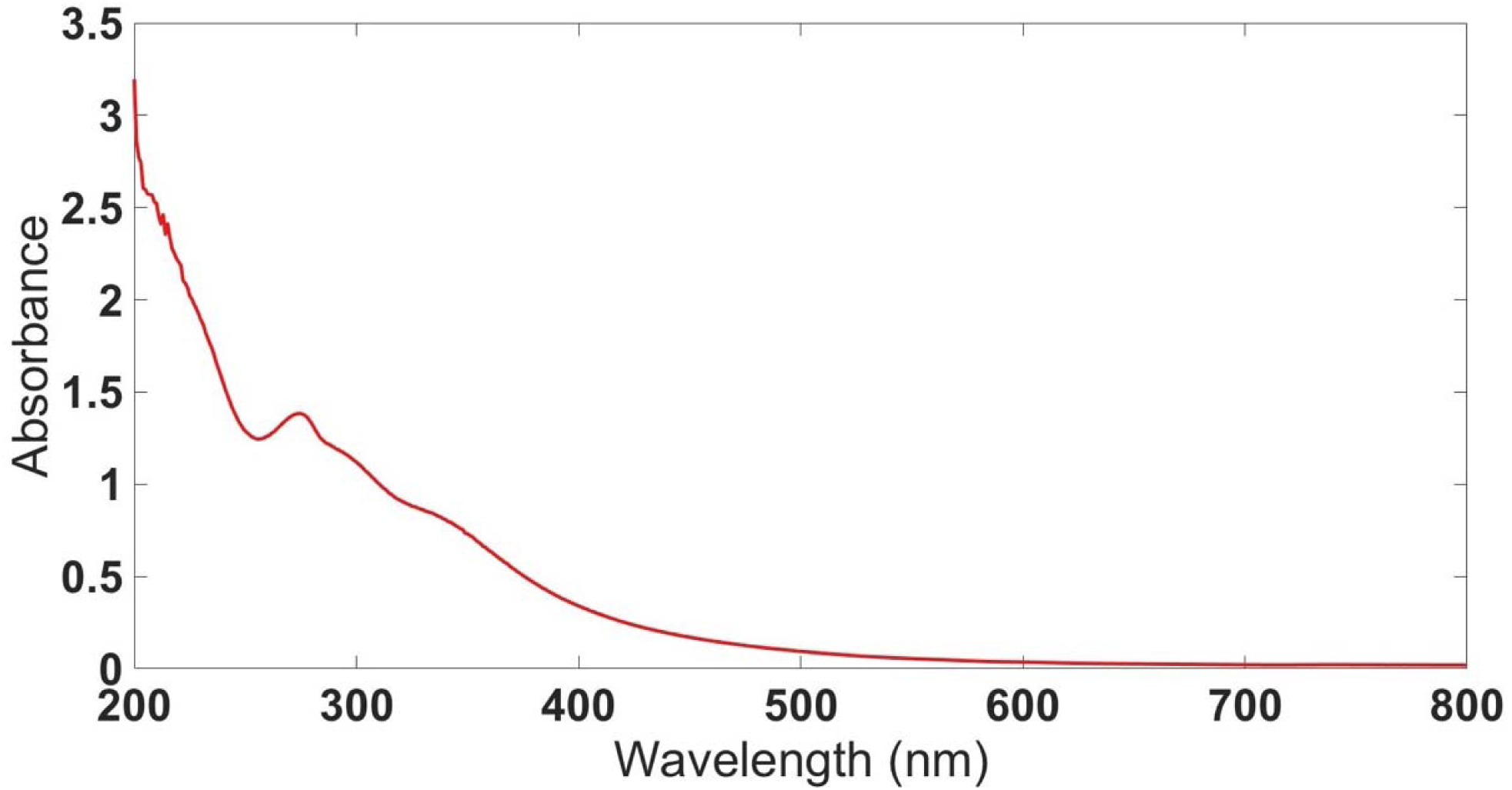
UV-visible absorption spectra for the PEG-CDs in ddH_2_O. The spectrum was taken at a concentration of 1.0 mg/ml in a 3mm path length quartz cell.

### Fluorescence Spectroscopy

Below is the 2D excitation-emission correlation fluorescence spectrum for the PEG-CDs. All fluorescence measurements were recorded using a Perkin-Elmer LS45 spectrometer. All spectra were acquired at 1.0 mg/ml in ddH_2_O in a 3mm path length quartz cell.

**Figure S4:**
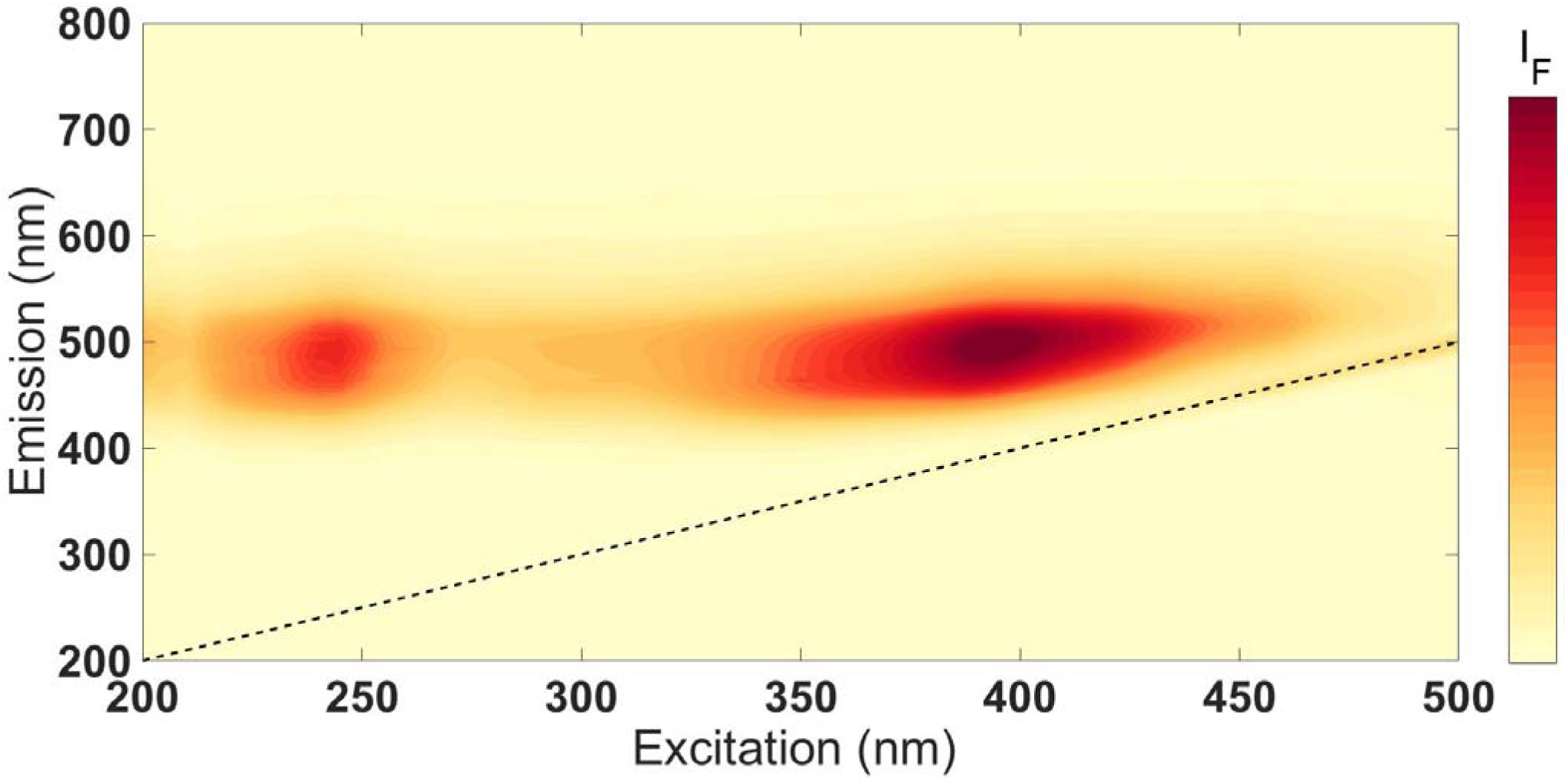
Two-dimensional excitation-emission correlation fluorescence spectrum of the PEG-CDs, I_F_ is the intensity of fluorescence and the black dashed line is the excitation laser.

### Dynamic light scattering (DLS)

Dynamic light scattering was performed on a Malvern Zetasizer nano at 25°C in water.

**Figure S5:**
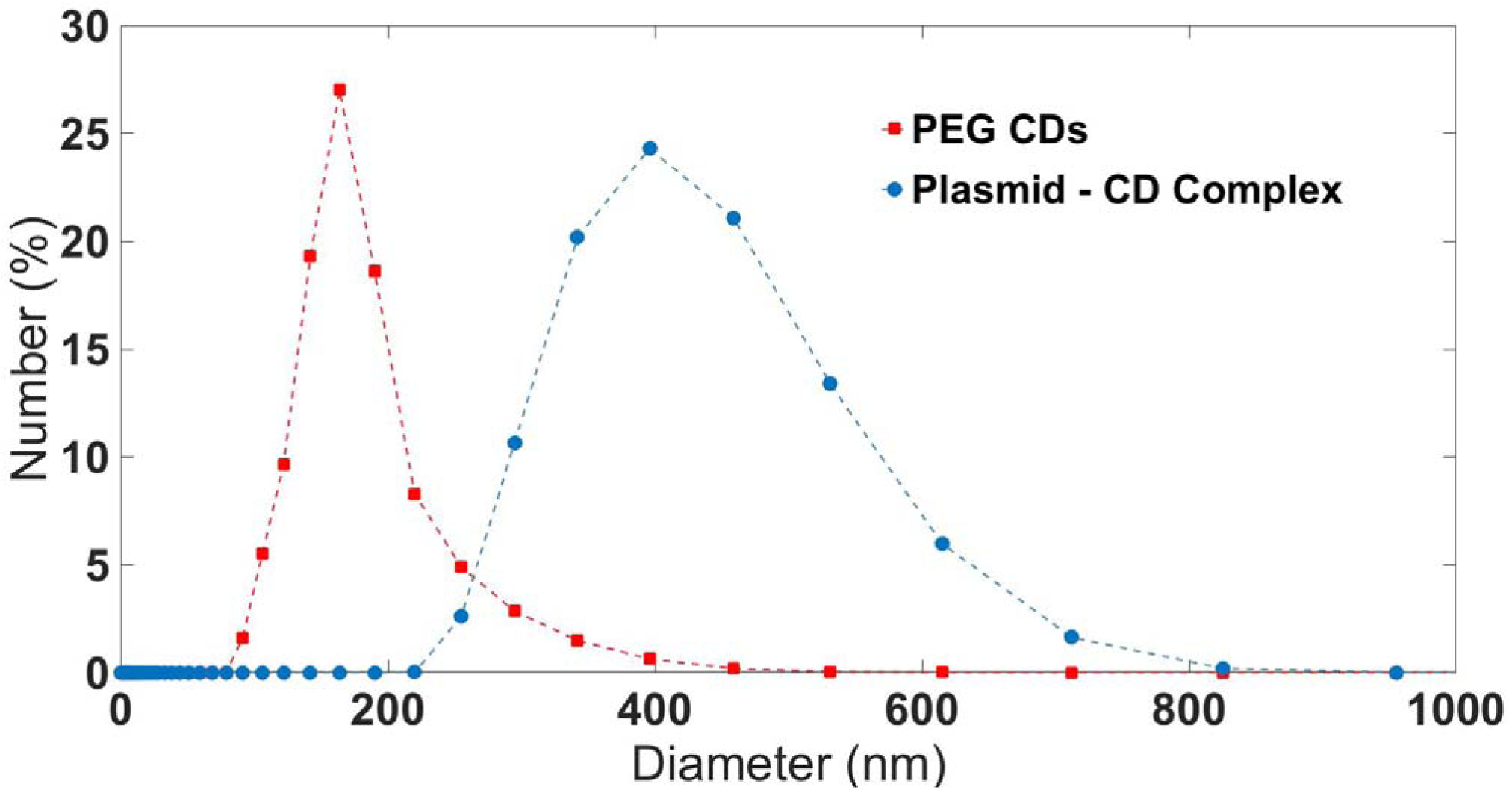
DLS of the PEG-CDs and the Plasmid-CD complexes.

### CDs do not cause growth defects

CD application by multiple methods to sorghum seedlings does not cause any obvious toxicity as no detrimental effects to growth are seen, even after multiple rounds of application (Fig. S6).

**Figure S6.**
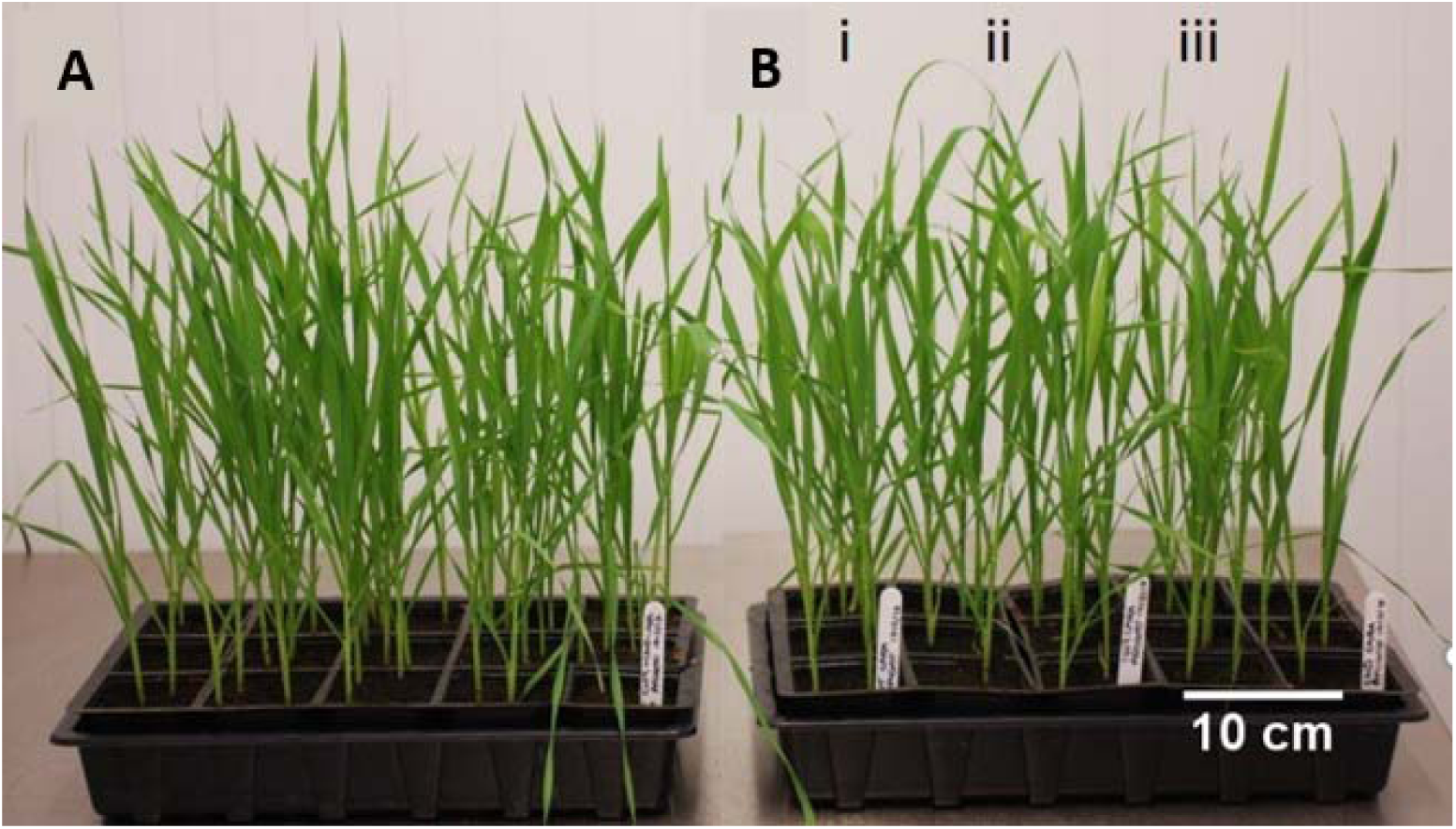
CD application is non-toxic. **(A)** Sorghum seedlings treated with CD-DNA via spraying **(B)**; i). Untreated Sorghum, ii). Sorghum control treated with DNA, no CDs, iii). Sorghum treated with CDs, no DNA. Each treatment was applied twice a day for five days.

### CD application does not affect photosynthesis

Spray on application of CDs (no DNA attached) to *Sorghum bicolor* did not statistically significantly affect the efficiency of photosynthesis at any PAR level (measured as ϕPSII), as determined by one-way ANOVA and Dunnett’s multiple comparison’s tests (Fig. S7).

The Maxi version of the IMAGING-PAM M-Series (Walz) was used to measure ϕPSII of CD treated *Sorghum bicolor*. Plants were left in the dark to acclimate for 1 hour before measurements were taken. Measurement settings were left as default. The Maxi-PAM measured the ϕPSII (operating efficiency of photosystem II. Results were exported for use in statistical analysis using the ImagingWinGigE V2.47+ (Walz) programme.

Matlab (Mathworks®) version 9.4.0.813654 (R2018a) was used to graph PAR and ϕPSII, and perform one-way ANOVA and Dunnett’s multiple comparison’s tests.

**Figure S7.**
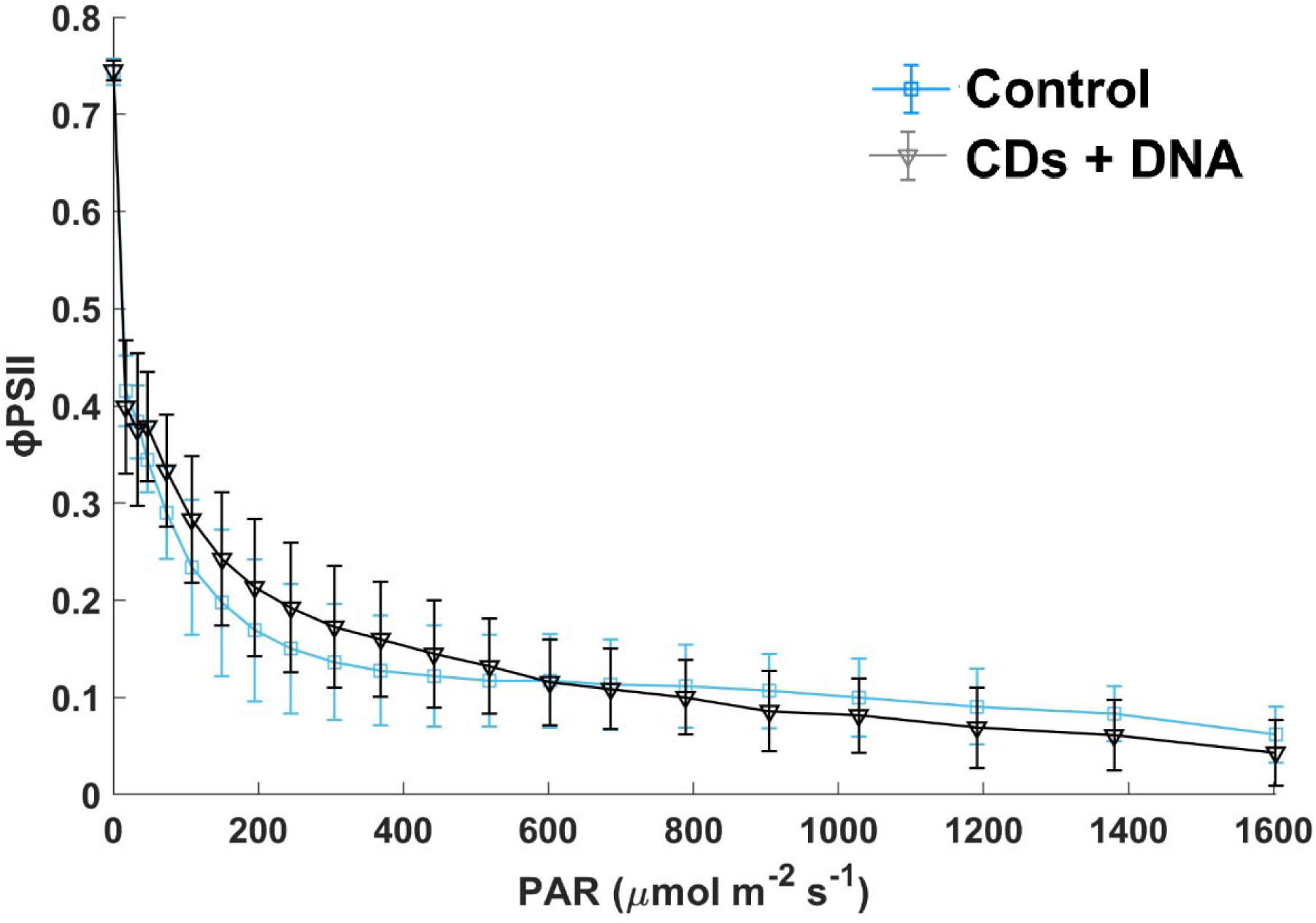
CD application has no significant effect on ϕPSII. Error bars indicate standard deviation of the mean.

### Sequencing confirms gene editing

DNA was extracted from wheat treated with CDs and our plasmid, and sequenced (Edwards Lab, UoB). The Multiple Sequence Alignment (Fig. S8) was created using T-Coffee^41^ and BoxShade.

**Figure S8.**
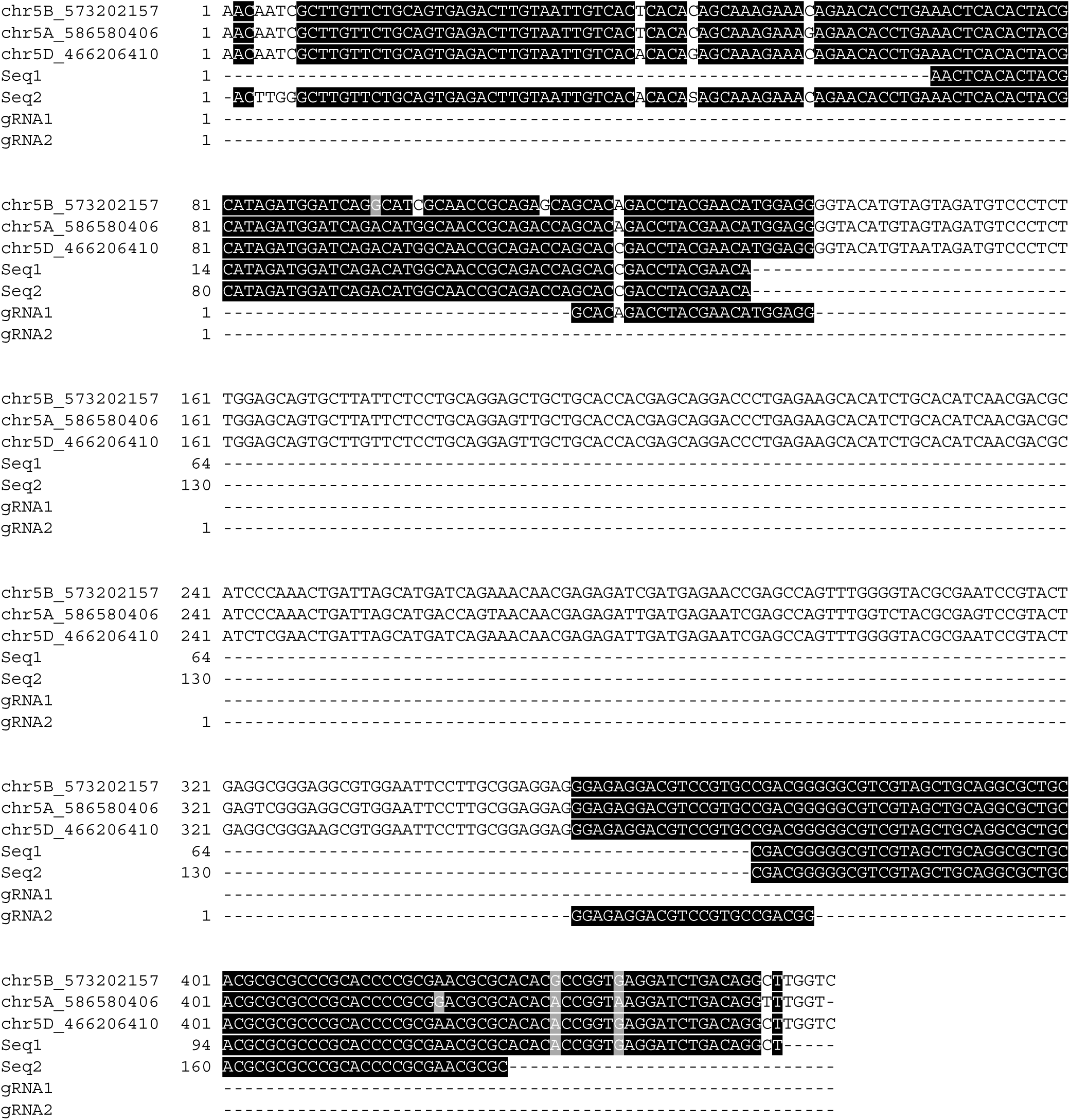
DNA sequencing shows gene edits in wheat. Note, wheat is hexaploid for the SPO11 gene. Image created using T-Coffee and Boxshade. Seq1 and Seq2 are forward and reverse Sanger sequencing reads from the same edit (Fig. 4.)

### Controls for PCR gels

Appropriate controls were performed for the gel shown in Fig. 4.

**Figure S9.**
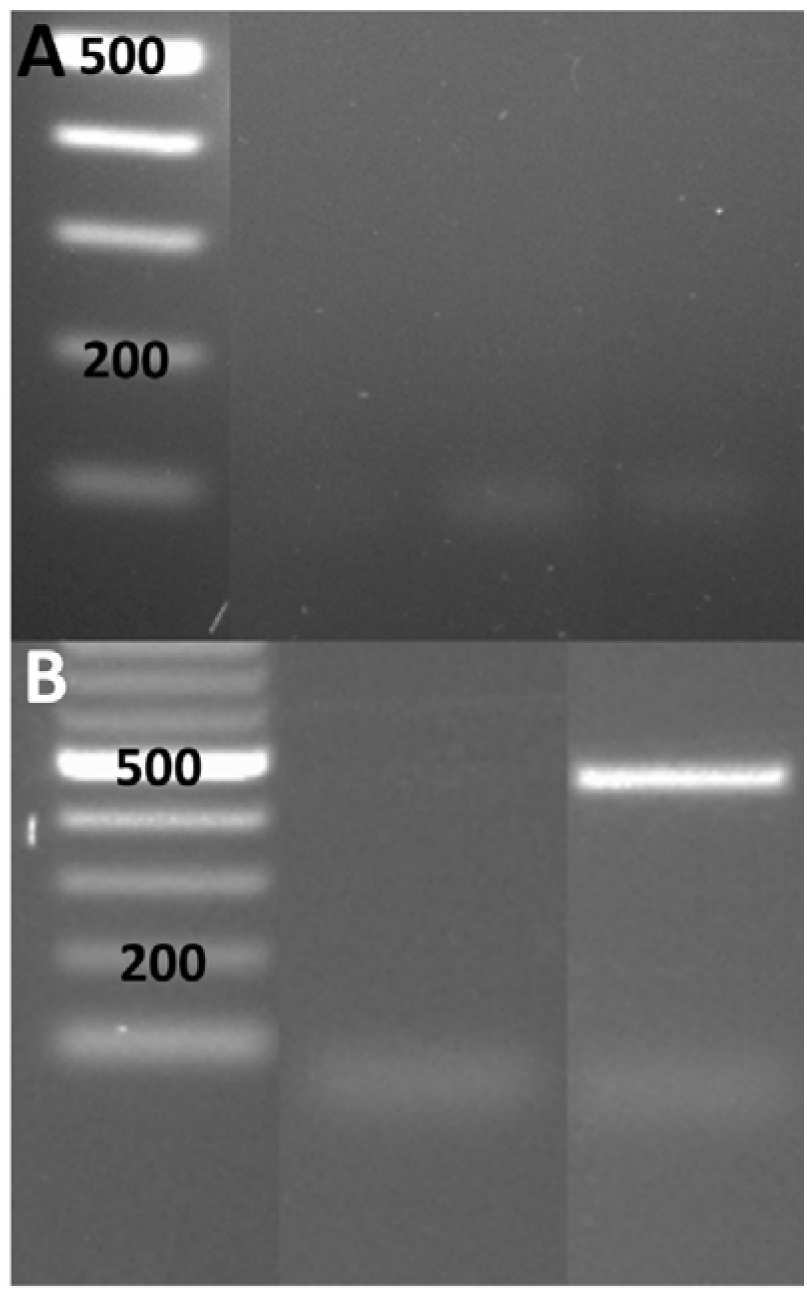
Gene editing controls. **(A), (B)** Controls for gene editing resulting in bands at c.230bp, obtained by **(A)**; CD sprayed leaf material, and **(B)** standard PEG-based transformation of protoplasts, as in Fig. 4. **(A)**, Lane 1, MW ladder (bp), Lane 2 (non-contiguous), full negative control, DNA extraction, restriction digest, and PCR reagents, Lane 3, negative control, restriction digest and PCR reagents, Lane 5 (non-contiguous), negative control, PCR reagents. **(B)**, Lane 1, MW ladder (bp), Lane 2 (non-contiguous), negative control, PCR reagents, Lane 3, positive control for plasmid with just GFP expression.

### Plasmid map

The structure of the pCas9-GFP plasmid is reported in Zhang *et al*., 2019^25^, but for ease of understanding a plasmid map is also provided here (Fig. S10).

**Figure S10.**
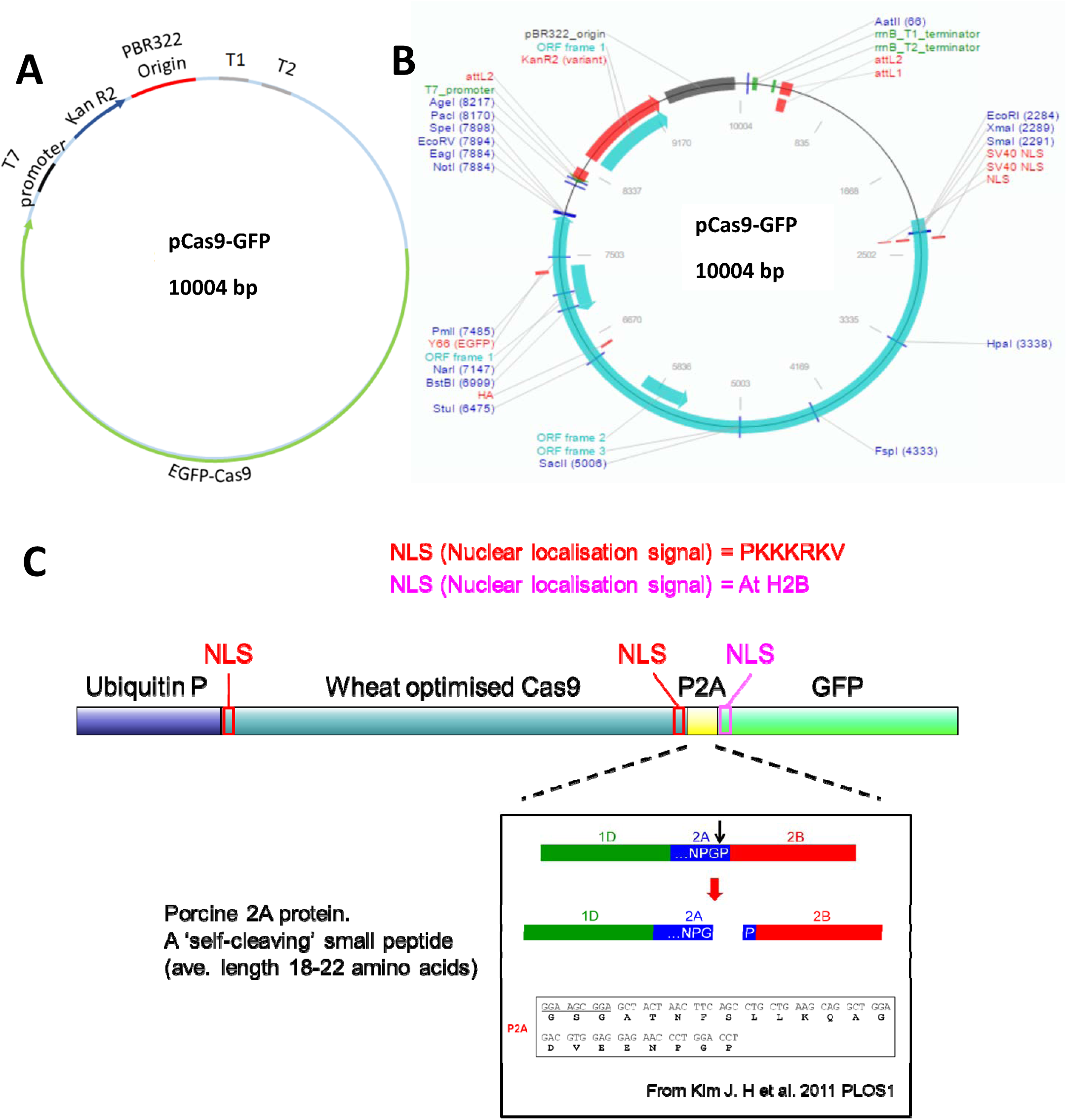
pCas9-GFP plasmid. **(A)**, Simplified pCas9-GFP plasmid map **(B)**, restriction map of pCas9-GFP, **(C)**, detailed map of the EGFP-Cas9 region **(A)** / ORF frame 3 region **(B)** of the pCas9-GFP plasmid.

