## Supplementary material for "A simple method for spray-on gene editing *in planta*": Safety guidance

### A simple spray on method for gene editing in planta – supporting safety information and guidance

*The below information is believed to be correct but does not purport to be all inclusive and shall be used only as a guide. The information in this document is based on the present state of our knowledge and is applicable to the products with regard to appropriate safety precautions. It does not represent any guarantee of the properties of the product. The authors of this paper and their institution's safety information shall not be held liable for any damage resulting from handling or from contact with the below products. Full material safety data sheets for substances that possess them must be consulted before work can begin, and the appropriate and relevant COSHH/health and safety forms should be filled in and checked and signed off by the appropriate persons at a given institution. The existence of this document is not endorsement for persons unfamiliar with any substances or equipment required by this protocol, especially the carbon nanomaterials, to begin synthesis and usage of them for plant transformation. Any persons wanting to begin this work should be fully trained in chemistry/biology safety protocols. Should you have any questions, please contact the Whitney or Galan labs for advice.*
*This document written by Katie Higginbottom with input from other authors of the paper.*

**Carbon Dot Synthesis**

Synthesis of Carbon Dot nanomaterials, functionalisation with PEGDA to form 1000MW PEG CDs.

| Substance | Form | Exposure routes | Hazards | First aid measures | PPE |
|---|---|---|---|---|---|
| **Glucosamine Hydrochloride** | Solid | Eyes, skin, inhalation, ingestion | May cause minor irritation | Flush eyes with water for 15 mins, flush skin with water for 15 mins, remove contaminated clothing and shower, if inhaled remove from exposure and move to fresh air, if not breathing, give artificial respiration, if ingested do NOT induce vomiting, rinse mouth and drink 2-4 cups of water | lab gloves, goggles, labcoat |
| **4,7,10-trioxa-1,13-tridecanediamine (TTDDA**) | Liquid | Eyes, skin, inhalation, ingestion | Burns, harmful to aquatic organisms | Flush eyes with water for 15 mins, flush skin with water for 15 mins, remove contaminated clothing and shower, if inhaled remove from exposure and move to fresh air, if not breathing, give artificial respiration, if ingested do NOT induce vomiting, | lab gloves, goggles, labcoat. Lab must have eyewash and safety shower. |

| | | | | | |
|---|---|---|---|---|---|
| | | | | rinse mouth and drink 2-4 cups of water. Do not let enter drains. | |
| **Methanol** | Liquid | Eyes, skin, inhalation, ingestion | Highly flammable liquid and vapour, toxic if swallowed, causes damage to organs | Flush eyes with water for 15 mins, flush skin with water for 15 mins, if inhaled remove from exposure and move to fresh air, if not breathing, give artificial respiration, if ingested call a poison centre or doctor/physician immediately, rinse mouth. Keep away from heat/open flames. In case of fire use CO2 powder/ alcohol resistant foam to extinguish. | Face shield, lab gloves, goggles, labcoat. Lab must have eyewash and safety shower. |
| **Tetrahydrofuran** | Liquid | Eyes, skin, inhalation, ingestion | Highly flammable liquid and vapour, harmful if swallowed, causes serious eye irritation, may cause respiratory irritation. Suspected of causing cancer. May form explosive peroxides. | Flush eyes with water for 15 mins, flush skin with water for 15 mins, if inhaled remove from exposure and move to fresh air, if not breathing, give artificial respiration, if ingested call a poison centre or doctor/physician if feeling unwell, rinse mouth. Keep away from heat/open flames. In case of fire use CO2 powder/ alcohol resistant foam/ water spray to extinguish. Do not let enter drains. | Face shield, lab gloves, goggles, labcoat. Lab must have eyewash and safety shower. |
| **CDI (1,1-carbonyldiimidazole)** | Solid | Eyes, skin, inhalation, ingestion | Harmful is swallowed, causes severe skin burns and eye damage, may damage the unborn child. | Flush eyes with water for 15 mins, flush skin with water for 15 mins, remove contaminated clothing and shower, if inhaled remove from exposure and move to fresh air, if not breathing, give artificial respiration, if | Face shield, lab gloves, goggles, labcoat. Lab must have eyewash and safety shower. |

| | | | | | |
|---|---|---|---|---|---|
| | | | | ingested do NOT induce vomiting, rinse mouth and drink 2-4 cups of water. | |
| **Polyethyleneglycol diamine (PEGDA)** | Solid | Eyes, skin, inhalation, ingestion | Not hazardous | Flush eyes with water for 15 mins, flush skin with water for 15 mins, remove contaminated clothing and shower, if inhaled remove from exposure and move to fresh air, if not breathing, give artificial respiration, if ingested do NOT induce vomiting, rinse mouth and drink 2-4 cups of water. | lab gloves, goggles, labcoat |
| **Carbon Dots (CDs)** 1000MW PEG CDs. | In solution. Will not dry out to form a powder. | Eyes, skin, inhalation, ingestion | The risks arising from carbon dots have not been fully characterised. They may be absorbed through the skin and may possibly be toxic if ingested. They are therefore treated as if harmful or potentially toxic. However, carbon dots have shown no cyto toxicity in mice and rats (Nanoscale Res Lett. 2013; 8(1): 122), and have shown non-toxicity in HeLa (human cervical) and MDA-MB-231 (human breast) cancer cells (Nanoscale 2016; 8(44): 11). | Flush eyes with water for 15 mins, flush skin with water for 15 mins, remove contaminated clothing and shower, if inhaled remove from exposure and move to fresh air, if not breathing, give artificial respiration, if ingested do NOT induce vomiting, rinse mouth and drink 2-4 cups of water. Do not let enter drains. | Face shield, lab gloves, goggles, labcoat. Lab must have eyewash and safety shower. Change nitrile lab gloves every 10 minutes. All carbon dot handling should be conducted inside a cat 2 fume hood with the ventilation on and glass pane pulled as far down as possible. Carbon dots should be in sealed vessel when outside of fume hood. |

**Carbon Dot based plant transformation**

Usage of Carbon Dots as a vehicle to carry plasmid DNA into plant cells for plant transformation.

| Substance | Form | Exposure routes | Hazards | First aid measures | PPE |
|---|---|---|---|---|---|
| **TE buffer** | Liquid | Eyes, skin, inhalation, ingestion | Not hazardous | Flush eyes with water for 15 mins, flush skin with water for 15 mins, remove contaminated clothing and shower, if inhaled remove from exposure and move to fresh air, if not breathing, give artificial respiration, if ingested do NOT induce vomiting, rinse mouth and drink 2-4 cups of water. | lab gloves, goggles, labcoat |
| **Plasmid DNA** | In solution | Eyes, skin, inhalation, ingestion | May cause slight eye irritation | Flush eyes with water for 15 mins, flush skin with water for 15 mins, remove contaminated clothing and shower, if inhaled remove from exposure and move to fresh air, if not breathing, give artificial respiration, if ingested do NOT induce vomiting, rinse mouth and drink 2-4 cups of water. | lab gloves, goggles, labcoat |
| **Carbon Dots (CDs)** 1000MW PEG CDs. | In solution. Will not dry out to form a powder. | Eyes, skin, inhalation, ingestion | The risks arising from carbon dots have not been fully characterised. They may be absorbed through the skin and may possibly be toxic if ingested. They are therefore treated as if harmful or potentially toxic. However, carbon dots have shown no cyto toxicity in mice and rats (Nanoscale Res Lett. 2013; 8(1): 122), and have shown non-toxicity in HeLa | Flush eyes with water for 15 mins, flush skin with water for 15 mins, remove contaminated clothing and shower, if inhaled remove from exposure and move to fresh air, if not breathing, give artificial respiration, if ingested do NOT induce vomiting, rinse mouth and drink 2-4 cups of water. Do not let enter drains. | Face shield, lab gloves, goggles, labcoat. Lab must have eyewash and safety shower. Change nitrile lab gloves every 10 minutes. All carbon dot handling should be conducted inside a cat 2 fume hood with the |

| | | | | | |
|---|---|---|---|---|---|
| | | | (human cervical) and MDA-MB-231 (human breast) cancer cells (Nanoscale 2016; 8(44): 11). | | ventilation on and glass pane pulled as far down as possible. Carbon dots should be in sealed vessel when outside of fume hood. |
| **Carbon Dots (CDs) 1000MW PEG CDs + DNA nanocomplexes** | In solution. Will not dry out to form a powder. | Eyes, skin, inhalation, ingestion | The risks arising from carbon dots have not been fully characterised. They may be absorbed through the skin and may possibly be toxic if ingested. They are therefore treated as if harmful or potentially toxic. However, carbon dots have shown no cyto toxicity in mice and rats (Nanoscale Res Lett. 2013; 8(1): 122), and have shown non-toxicity in HeLa (human cervical) and MDA-MB-231 (human breast) cancer cells (Nanoscale 2016; 8(44): 11). | Flush eyes with water for 15 mins, flush skin with water for 15 mins, remove contaminated clothing and shower, if inhaled remove from exposure and move to fresh air, if not breathing, give artificial respiration, if ingested do NOT induce vomiting, rinse mouth and drink 2-4 cups of water. Do not let enter drains. | Face shield, lab gloves, goggles, labcoat. Lab must have eyewash and safety shower. Change nitrile lab gloves every 10 minutes. All carbon dot handling should be conducted inside a cat 2 fume hood with the ventilation on and glass pane pulled as far down as possible. Carbon dots should be in sealed vessel when outside of fume hood. CD/DNA nanocomplex spraying should only ever occur in a dedicated cat 2 fume hood that is clear of any clutter or other items. |

**Extra Carbon Dot information**

- All Carbon Dot waste (pipette tips, empty spray bottles, wipes used to clean up spills, plant tissue) should be contained within biohazard bags to prevent release and disposed of by incineration.
- If CDs are spilled, soak a towel in 70% ethanol and mop up the spill. Then, wipe the area again with another towel. Both towels are waste for incineration.
- All equipment should be cleaned in 70% ethanol.
- All containers containing CDs must be clearly marked as containing "carbon nanomaterials, PEG 1000MW carbon dots", and marked as hazardous. Waste must also be labelled as such. "CD" is insufficiently informative to be used on labelling.
- Questions/queries about these carbon nanomaterials should be directed to the Galan research group.
- Questions/queries about the transformation protocol/gene editing should be directed to the Whitney and Edwards research groups as appropriate.

**Spray protocol specific**

- Carbon dot transformation spraying should only occur in a dedicated cat 2 fume hood – this prevents exposure to others, will prevent exposure to other lab plants/materials, and minimise the number of areas to be kept clean of CD waste.
- Spray plants from approx. 10cm away. Aim downwards. Rotate the plant rather than moving the spray around. Do not spray towards fume hood glass. When spraying, ensure glass pane is pulled down as far as possible, leaving as small a gap as possible for your hands and wrists.
- Plants brought from growing rooms to cat 2 fume hood for spraying should have an extra tray/box as a secondary level of containment that should not enter the fume hood. Ideally should shut securely. As below.

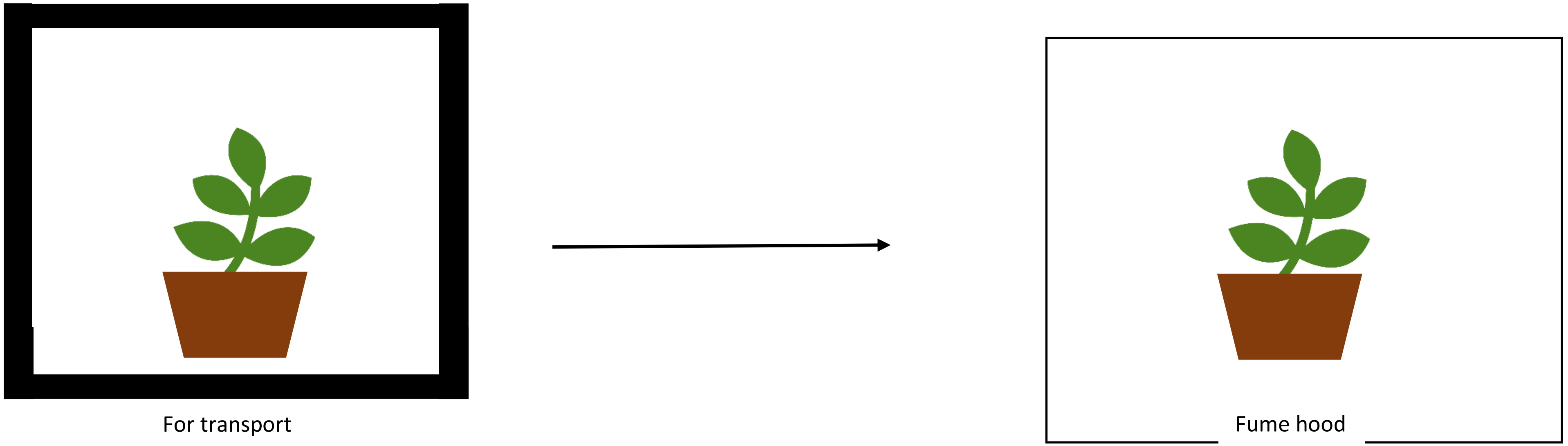

- Once sprayed, plants should be left to dry completely before removal from the cat 2 fume hood. This may take upwards of 20 minutes.
- CD treated plants should be kept in isolation from non CD treated plants.
- CD treated plants should have a water drainage system separate from non CD treated plants. This waste must be disposed of as above.
- Do not transport CD materials or treated experimental substances through public areas such as lobbies or any non-lab corridor.

**Gene Editing**

- gRNAs for Cas9 gene editing should be designed so as to minimise the amount of off target effects and to be as species specific as possible, should contact to other organisms occur.
- GMO/gene edited plants should be kept isolated in a dedicated GMO plant growth room.
- Check recent literature for up to date safety guidance and protocols for the use of CRISPR-Cas9 machinery.
